# Inverse association between serum vitamin B_12_ level and abundance of potential B_12_-producing gut microbes in Indian children

**DOI:** 10.1101/2025.04.21.649764

**Authors:** Nisha Chandel, Priyansh Patel, Pramod R. Somvanshi, Anil Kumar Verma, Vivek Thakur

## Abstract

**Background:** The human gut microbiome is a natural source of essential micronutrients like B-vitamins, which are utilized by both the host and other community members. The prevalence and abundance of known B-vitamin producers and B-vitamin biosynthesis pathways have already been reported in gut microbiome cohorts of a few countries including India.

**Objective:** To test whether the presence of B-vitamin producers/biosynthetic pathways translates into serum B-vitamin levels, taking B_12_ as a case example.

**Methods:** Fecal samples were collected from non-deficient (serum B_12_ level > 210 pg/mL, n=29) and B_12_ deficient (serum B_12_ level < 210 pg/mL, n=30) children from a tribal region of central India. Whole metagenomic DNA was extracted, sequenced, and analyzed for taxonomic profiling and diversity comparisons. Differentially abundant taxa between two groups were identified. The prevalence and abundance of potential B_12_ producers were compared, and their association with serum B_12_ level was established.

**Results:** A comparison of *within-sample* diversity between the two groups didn’t show any difference; however, *between-sample* diversity was significantly less in the B_12_ deficient group. Differential abundance testing also showed different microbiome structure in the B_12_ deficient group, where an increased abundance of B_12_ transporter-carrying *Bacteroides thetaiotaomicron*, a few pathogenic species, and ten known B_12_ producers was observed. Potential B_12_ producers were also significantly prevalent and abundant in the deficient group. Their cumulative abundance was also significantly higher in the deficient group and showed a negative association with serum B _12_ levels.

**Conclusion:** A higher abundance of potential B_12_ producers in the deficient group suggested an adaptive mechanism by the gut microbiome to meet the community’s B_12_ requirements, by selectively promoting the growth of B_12_ producers.

## Introduction

Cobalamin (Vitamin B_12_) is one of the structurally most complex biomolecules, exclusively synthesized by bacteria and a few archaea [1, 2]. It is needed as a co-factor only for two enzymes in humans, namely, cytosolic Methionine synthase (MetH) and mitochondrial L-Methyl-malonyl-CoA mutase (MCM) [3]. Being an essential micronutrient, it must be acquired by ingestion [1]. Its deficiency impairs DNA synthesis, and causes erythroblast apoptosis, resulting in anemia (reviewed by Koury and Ponka; 2004) [4]. B_12_ depleted diet, malabsorption, and defects in the cellular delivery and uptake system are the causes of deficiency, as reviewed in multiple studies [5–7]. The recommended dietary intake (RDI) of vitamin B_12_ (Vit. B_12_) ranges from 0.9 to 2.4 micrograms from infants to adults, which can be sourced from an animal-based diet, and fortified foods [8]. Ingestion of such foods brings Vit. B_12_ to the small intestine, where it binds to an intrinsic factor (IF) released from the stomach. This IF-B_12_ complex gets absorbed in the small intestine. The unabsorbed B_12_ is utilized by bacteria in both the small and large intestine (see the review articles) [7, 9].

Biosynthesis of Vit. B_12_ is one of the essential functions of the gut bacterial community [10, 11]. However, approximately 25% of sequenced gut bacterial genomes had the genetic potential to synthesize corrinoids (cobalamin and related cofactors), whereas 80% were predicted to use them [12], indicating that the majority of the community members depend on B_12_ producers and unabsorbed dietary B_12_ for their requirement.

Vit. B_12_ modulates the gut community structure, as it is best known for its role as a cofactor and gene expression regulator in bacteria [13]. Different gut bacterial species require distinct corrinoids, and they obtain specific ones by corrinoid remodeling, which limits their availability to the host [13, 14]. Several possible competitive and symbiotic interactions within the gut community indicated the co-evolution of community members, which possibly has also benefited the vitamin homeostasis of the host [11].

Earlier, we reported the biosynthetic potential of gut microbes for all B vitamins in Indian cohorts [10] and estimated a slightly higher contribution of such microbes to RDI than a previous report [11]. On the contrary, a contemporary country-wide nutritional survey (Comprehensive National Nutrition Survey, CNNS-2018) reported a worrying level of prevalence of Vit. B_12_ deficiency in the Indian population [15]. This made us wonder if i) a high abundance of B _12_ producers or B_12_ biosynthesis pathways in the gut gets translated into host serum B_12_ levels, and ii) serum B_12_ deficiency shows any association with the gut community structure. The current study aims to answer these questions by analyzing fecal metagenome data from B_12_ deficient (n=30) and non-deficient (n=29) children from the Balaghat district of Madhya Pradesh, India, where we assumed that fecal metagenome is a proxy of the gut metagenome and tested the association of host serum B_12_ level with potential gut B_12_ producers (Figure 1A).

**Figure 1.**
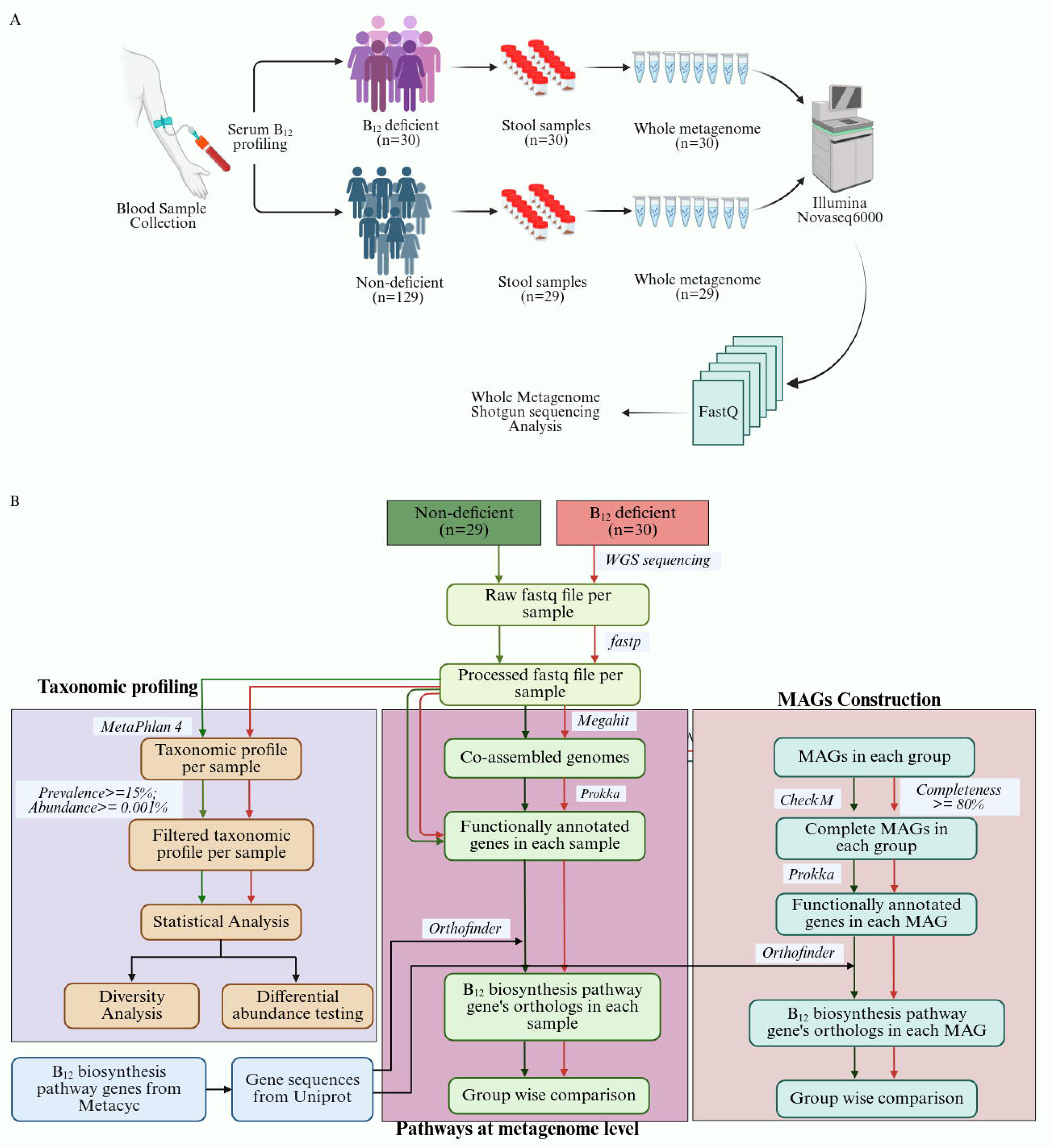
Methodology. **A** sampling and **B** computational analysis. BioRender software was used to create the figure under an academic license.

## Materials and Methods

### Sample Collection

Sampling sites included the Nutrition Rehabilitation Centre (NRC), Anganwadi centres (part of the Integrated Child Development Services program), and the Government Primary School of wards 3, 4, and 9 of Baihar village in the Balaghat district of Madhya Pradesh state, India. Children who hadn’t taken antibiotics for the last month and didn’t have diarrhoea, constipation, other gastrointestinal diseases, or any other disease were included in this study. Metadata about all subjects such as age, gender, BMI, diet, and antibiotics or supplements usage, was gathered using a questionnaire and consent form, which was approved by the Institutional Ethics Committee (UH/IEC/2022/389), University of Hyderabad, India and ICMR-National Institute of Research in Tribal Health, India (Table S1). The basic summary of the data for the two groups is provided in Table 2. The fecal sample collection was done from January 2023 to March 2023, involving children with ages ranging between 2-12 years as follows: serum B_12_ profiling of 158 children was first done at Thyrocare Technologies Ltd. (Mumbai) using Chemi Luminescent Microparticle Intrinsic factor Immunoassay (CMIA; Intra assay (%CV): 5.6, Inter assay (%CV): 6.8, Sensitivity: 125 pg/mL) or Competitive Chemi Luminescent Immuno Assay (CLIA; Intra assay (%CV): 5.0%, Inter assay (%CV): 9.2 %; Sensitivity: 45 pg/ml) methods, and they were classified into non-deficient (>210 pg/ml; n=118) and B_12_ deficient groups (<210 pg/ml; n=40). After obtaining 40 B_12_ deficient cases, further screening of serum B_12_ profiling was stopped as it was an exploratory study. The fecal samples, however, could be obtained only for 30 of them. To make both groups comparable, 29 non-deficient children were randomly selected for fecal sampling. About 30 participants per group is a common starting point in exploratory research due to its ability to balance statistical power with practical feasibility, its suitability for detecting small-to-medium effects in high-dimensional microbiome data, and its alignment with power analysis estimates for achieving a reasonable likelihood of detecting true effects in small-to-medium-sized studies. The fecal samples were collected in sterile stool containers and stored at 0°C within 1 to 5 hours and later were transported in dry ice to miBiome Therapeutics LLP Mumbai, India, for DNA extraction and sequencing (Figure 1A).

**Table 1.** List of the B_12_ biosynthesis pathways or sub-pathways examined in this study, number of MAGs from each group having one of the pathways, and *p-value* of comparison of their proportion between groups.

| Pathway/Subpathway name | Short name | MetaCyc ID | Non-deficient | B<sub>12</sub> Deficient | *p*-value |
| --- | --- | --- | --- | --- | --- |
| Adenosylcobalamin biosynthesis (anaerobic) | anaerobic | PWY-5507 | 27 | 46 | 0.11 |
| AdoCbl biosynthesis from adenosylcobinamide-GDP I | GDP-1 | PWY-5509 | 63 | 85 | 0.3505 |
| AdoCbl biosynthesis from adenosylcobinamide-GDP II | GDP-II | PWY-7975 | 1 | 1 | – |
| AdoCbl salvage from cobalamin | Cobalamin salvage | PWY-6268 | 6 | 5 | 1 |
| Superpathway of AdoCbl salvage from cobinamide | Cobinamide salvage | COBALSYN-PWY | 44 | 67 | 0.139 |
| **Total MAGs** |  |  | **135** | **161** |  |

**Table 2.** Summary of the metadata collected from samples.

|  | **Non-deficient (n=29)** | **B<sub>12</sub> deficient (n=30)** |
| --- | --- | --- |
| **Serum B<sub>12</sub> level**<br>(pg/ml) | 392.83 (±156.73) | 161.63 (±30.80) |
| **Age** in years (mean±<br>(sd)) | 6.4 (±2.6) | 7.8 (±2.6) |
| **Weight** in kg (mean±<br>(sd)) | 16.3 (±5.0) | 18.8 (±4.8) |
| **BMI** (mean±(sd)) | 13.1 (±2.3) | 13.9 (±2.1) |
| **Gender** | Female=16, Male=13 | Female=21, Male=9 |
| **Diet** | Nonvegetarian=24, Vegetarian=5 | Nonvegetarian=30 |
| **Antibiotics** | No | One subject took antibiotics<br>2 months before sampling |

### Fecal Metagenomic DNA extraction and quantification

Whole metagenomic DNA from all 59 samples was isolated using the QIAamp PowerFecal Pro DNA Kit according to the manufacturer’s instructions. The DNA samples were quantified on NanoDrop One and Qubit using water and a standard as a control, respectively. DNA quality was estimated by agarose gel electrophoresis.

### Library Preparation and Sequencing

Libraries were prepared using QIAseq® FX DNA library kit from Qiagen. Briefly, 150 ng of DNA was used for fragmentation and end repair, followed by adapter ligation. Excess adapters were removed using AMPure XP purification beads, followed by PCR enrichment of adapter-ligated DNA and clean-up of library products as instructed by the manufacturer. The cleaned libraries were quantitated on a Qubit fluorometer, and appropriate dilutions were loaded on HS D1000 screen tape to determine the size range of the fragments and the average library size. Eventually, paired-end sequencing, with reads of size 150 bases each, was done using the Illumina *Novaseq6000* platform.

### Pre-processing of metagenomic reads

The quality of sequencing data was checked using FastQC v0.12.1 [16] and MultiQC v1.22.1 [17]. Since all reads had an average *Phred-*score > 25, only ambiguous bases and adapters were removed using *fastp* [18]. Further, host contamination in the filtered reads was removed by aligning reads to the human genome (GRCh38) using Bowtie2 v2.5.4 [19], and unaligned reads were extracted using SAMtools v1.20 [20]. The processed reads were used for downstream analysis (Figure 1B).

### Taxonomy profiling and species filtration

The compositional profiling of each sample was done using MetaPhlAn 4.0.6 (Metagenomic Phylogenetic Analysis) [21]. To further reduce the risk of contamination or alignment artifacts, species with very low mean relative abundance (<= 0.001%) or prevalence (<= 15%) were removed (Figure 1B).

### Diversity analysis

The filtered compositional profiles were used to measure alpha and beta diversity between the two groups. The alpha diversity (or within-sample diversity) was measured by using the *vegan* package [22] using multiple measures such as species richness (*specnumber()* function), Shannon index (*diversity()*), and Simpson index (*diversity()*). The beta diversity (or between-sample diversity) analysis was first performed using a dimensionality reduction approach: Principal Coordinate Analysis (PCoA). It was done using the *cmdscale()* function on Bray-Curtis distance matrix. Alternatively, the beta diversity analysis was performed using pair-wise distance measures like Bray-Curtis, Jaccard distance, and Robust Aitchison distances; all were calculated using the *adonis()* function in the *vegan* package in R.

### Vit. B_12_ pathway analysis at the community (or microbiome) level

For Vit. B_12_ biosynthesis pathway analysis, filtered reads of samples from each group were co-assembled using the *de novo* assembly tool, MEGAHIT (v1.2.9) [23]. The contigs of each co-assembly were annotated using the Prokka tool (v1.14.5) [24]. Gene abundance in each sample was estimated by mapping filtered reads from the respective sample to the gene sequences of the corresponding co-assembly using the Bowtie2 tool (v2.5.4) [19], and the read counts were estimated by the Expectation-Maximization-based tool, RSEM (v1.3.3) [25] (Figure 1B).

To examine the presence of B_12_ biosynthesis pathway(s) in any sample, the reference genes of the known B_12_ biosynthesis pathways or subpathways in bacteria were first obtained from MetaCyc [26] (Table 1). Since the majority of the reference genes were from *Salmonella typhimurium* genome (RefSeq ID: GCF_000006945.2) [27], the genome annotation using Prokka tool [24] was used to obtain the protein sequences. The reference genes which were from other bacteria, namely, *Listeria innocua serovar 6a* (strain ATCC BAA-680 / CLIP 11262), *Escherichia coli (strain K12)*, *Priestia megaterium*, *Desulfuromonas acetoxidans* (strain DSM 684 / 11070), *Eubacterium callanderi*, their protein sequences were obtained from UniProt database [28], and appended to *S. Typhimurium* protein sequences. The orthologs of the appended set were identified among translated gene sequences of each co-assembly using the OrthoFinder tool (v2.5.5) [29] (Figure 1B).

The pathway abundance in each sample was estimated by taking the median value of normalized abundances of all genes of the pathway. The abundance of any gene was independent of the genomes to which they belonged.

### Vit. B_12_ pathway analysis at Metagenome-Assembled Genomes (MAGs) or species level

The above analysis was based on B_12_ biosynthesis through cooperativity at the community level. Additionally, the contribution from the individual MAGs with complete pathways was also considered for comparison between the groups. The contigs from each co-assembly were binned to obtain MAGs using the CONCOCT tool (v1.1.0) [30]. Their quality was assessed with CheckM2 tool (v1.0.2) [31], and MAGs with completeness <= 90 % were discarded. Complete MAGs were annotated using the Prokka tool (v1.14.5) [24], and their taxonomy was predicted using GTDB toolkit [32]. Orthologs of the B_12_ biosynthesis pathways gene in each MAG were identified by using the OrthoFinder tool (v2.5.5) [29] (Figure 1B). The mean contig depth (calculated by CONCOCT) belonging to a particular MAG was used as its abundance. The proportion of samples having a MAG was termed as its prevalence. Pathways with completeness <= 80% and present only in 5 MAGs or less than that were eliminated from the downstream analysis.

### Differential abundance testing and other statistical Analyses

All statistical analyses were performed in R (version 4.2.2; www.r-project.org). The difference in beta diversity was statistically tested using the *adonis()* function in the vegan package, which uses PERMANOVA. Statistical testing for differentially abundant species was done using a negative binomial regression model-based *Wald* test implemented in DESeq2 [33] after controlling for the effect of confounders, namely batch, age, gender, diet, and BMI, which uses Benjamini-Hochberg procedure (FDR) for multiple hypothesis testing. The adjusted *p-value (p-adj) <= 0.05* was considered significant. The known B_12_ producers were inferred by comparing the differentially abundant species to a comprehensive list from our previous study [10].

All the statistical comparisons between groups were performed using the non-parametric (*Wilcoxon rank-sum)* test. The proportion of MAGs carrying B_12_ pathways (potential B_12_ producers) between the two groups was compared using the proportionality test function (*prop.test()*). The association between serum B_12_ levels and the cumulative abundance of B_12_ producers was assessed using a multiple linear regression model in R (*lm()*), controlling for potential confounders such as age, gender, diet, and BMI. All figures were generated using the *ggplot2* package [34] in R.

## Results

### The prevalence of B_12_ deficiency in the tribal children cohort was higher than the national/state average for school-age children

Serum B_12_ profiling of 158 children from the Balaghat district of central India showed a deficiency in ∼25% of samples (Supplemental Table 2). Compared to CNNS-2018 survey data, though the prevalence in the current study had a similar increasing trend with age (10%, 28%, and 39.3%, in pre-school, school-age and adolescent, respectively), however, the prevalence in the school-aged group was relatively higher (∼28%, n=90) (Supplemental Table 2).

From the total of 158 subjects, two groups of similar size were made (see methods) for further study: one with a normal B_12_ level (n=29, range: 212-810 pg/ml, median=365 pg/ml) and another with B_12_ deficiency (n=30, range: 82-198 pg/ml, median=169 pg/ml) (Figure 1A, Supplemental Figure 1A; Table 2; Supplemental Table 1). Whole metagenome shotgun sequencing of fecal samples of both the groups resulted in ∼ 25 million paired-end reads per sample (∼24 million after quality control) (Supplemental Figure 1B).

### Mixed trend for alpha (*within-sample*) diversity in the B_12_ deficient group

Taxonomic profiling results showed the presence of 1225 species (Supplemental Table 3), where most of the species belonged to phylum Bacillota, Bacteroidota, Actinomycetota, and Pseudomonadota (Figure 2A). It was also observed that ∼20% of the reads remained unclassified, which may be partly due to the uniqueness of the Indian gut microbiome (Figure 2A). Species present only in a few samples and with low abundance were filtered (see materials and methods), and the remaining 374 species were considered for downstream analysis. Within-sample diversity showed no significant difference in the B_12_-deficient group when estimated using metrics based on both the richness and evenness, such as the Shannon index (*p* = 0.11) and Simpson index (*p* = 0.10). However, it was significantly higher when estimated using only the species richness (*p* = 0.05) (Figure 2B and C; Supplemental Figure 2A).

**Figure 2.**
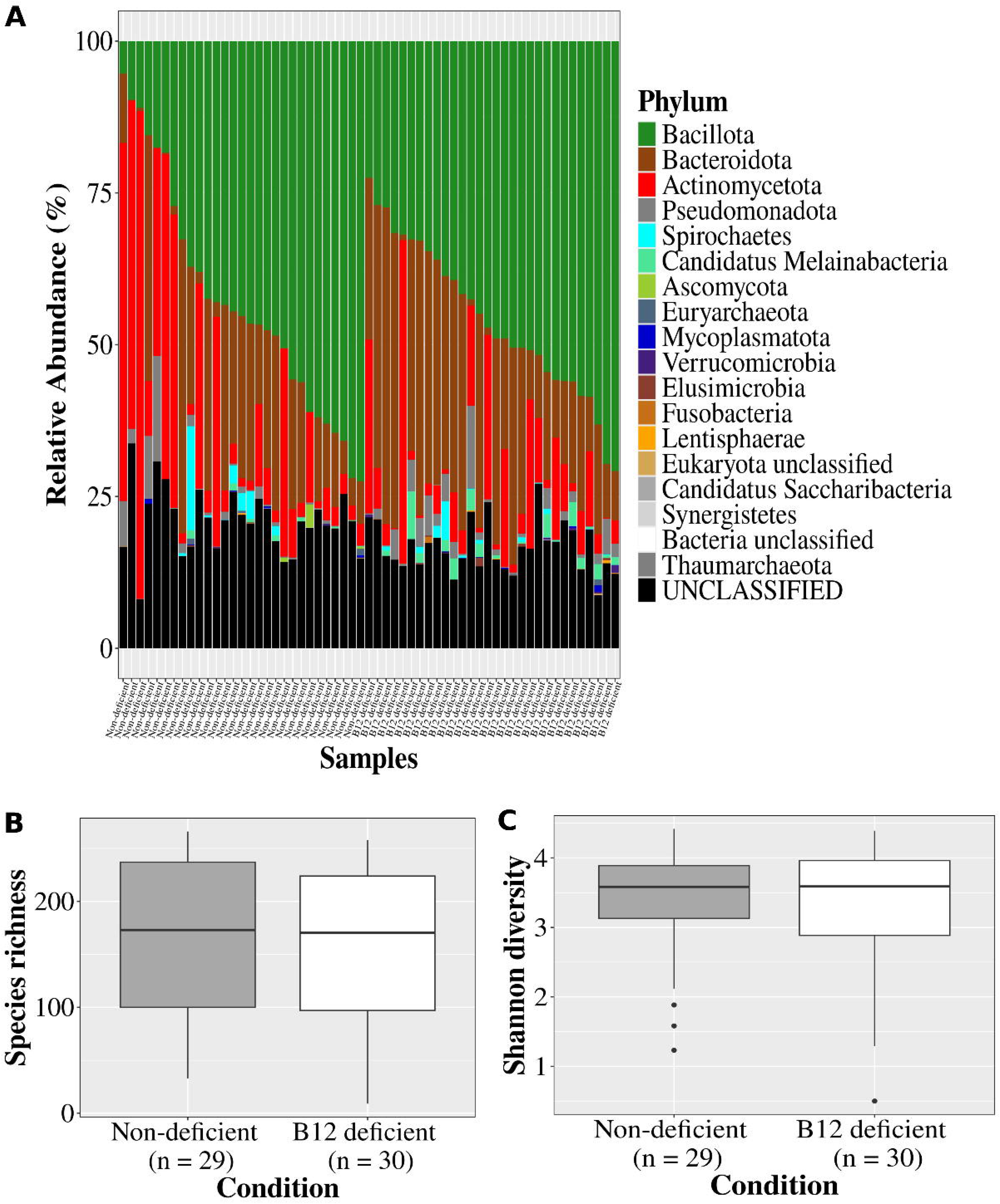
Alpha (within-sample) diversity trends. **A** Bar plot showing the relative abundance of detected phyla in the data. The Box plots represent within-sample diversity between Non-deficient and B_12_ deficient groups: **B** Richness (*p-value = 0.05*), and **C** Shannon diversity (*p-value = 0.11*).

### Significant reduction in beta (*between-sample*) diversity in the B_12_ deficient group indicated convergence

To examine if the microbiome composition was different in B_12_-deficient condition, PCoA using the Bray-Curtis dissimilarity index was done. B_12_ deficient samples clustered together, partially overlapping with the non-deficient ones along the first principal coordinate, which captured ∼24% of the variation (Figure 3A). The higher similarity (i.e., lower distance) among the B_12_ deficient samples compared to non-deficient ones was statistically confirmed using the PERMANOVA test (*p* = 0.004) (Figure 3B). Similar trends of lower beta diversity in the B_12_ deficient group were observed using other distance measures like the Jaccard distance (*p* = 0.005) and the Robust Aitchison distance (*p* = 0.07), further confirming different gut community structure in B_12_ deficient condition (Figure 3C; Supplemental Figure 2B).

**Figure 3.**
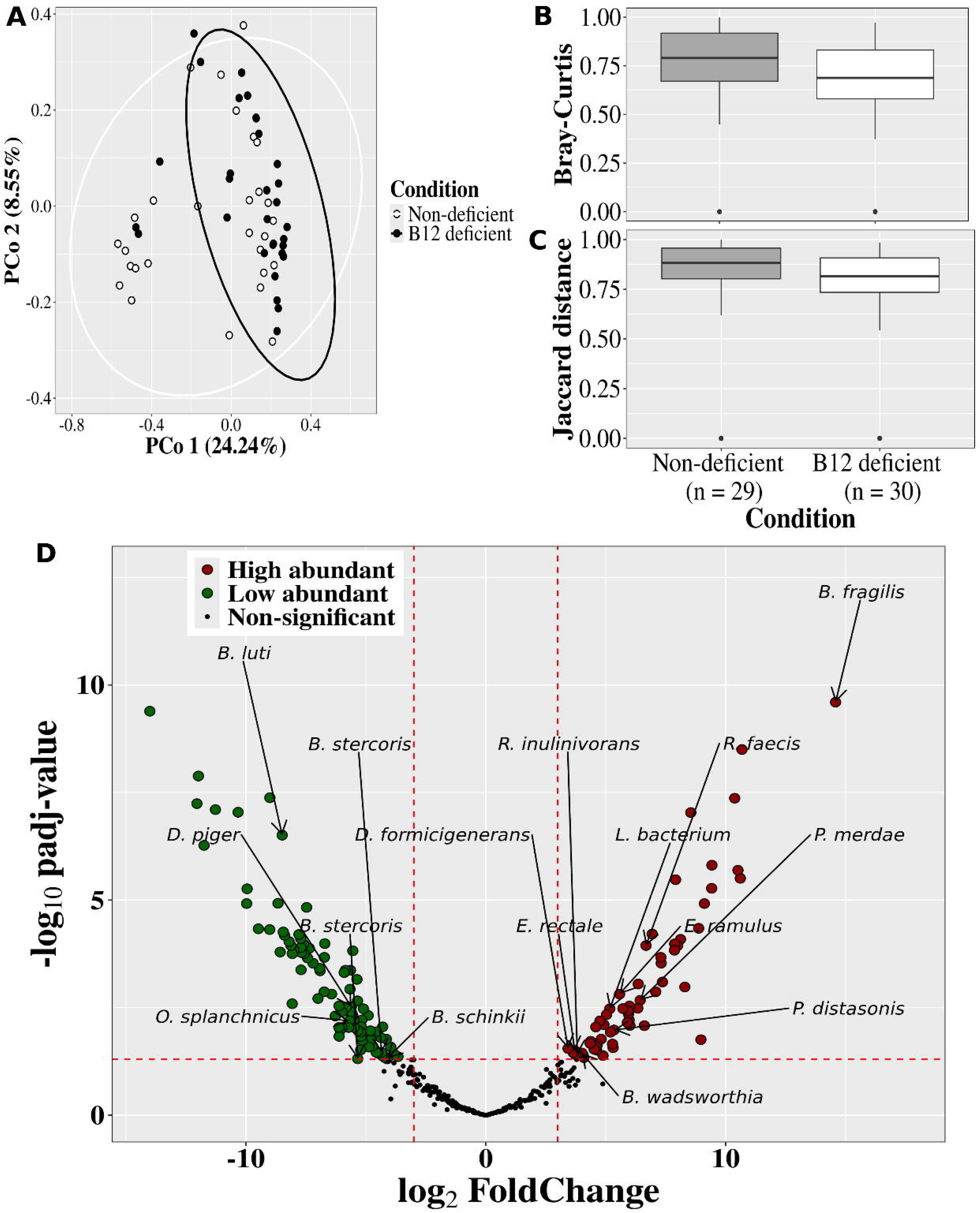
Beta (between-sample) diversity trends. **A** Principal coordinate analysis of Bray-curtis distances, **B** Bray-curtis dissimilarity index (*p-value = 0.004*), **C** Jaccard distance (*p-value = 0.005*), and **D** Volcano plot showing differentially abundant species between non-deficient and B_12_ deficient groups (*p-adj <= 0.05*). Differentially abundant known B_12_ producers have been highlighted.

### Ten *known* B_12_ producers were among sixty-two differentially more abundant species in the B_12_ deficient group

To find out differentially abundant species in any of the groups, their abundance was statistically compared. A total of 157 significantly abundant species, 62 were more abundant in the B_12_ deficient group (FDR = 0.05, log_2_ fold change >=3) (Figure 3D), which included the B_12_ transport system carrying species of genus *Bacteroides,* pathogenic species *Sutterella faecalis, Clostridiaceae unclassified SGB4771,* mucin degrader *Akkermansia muciniphila*, and others, which are listed in Supplemental Table 4 [35–38]. The non-deficient group had a higher abundance of lactate metabolizing species of genus *Veillonella*, *Streptococcus thermophilus*, plant-based diet associate *Prevotella stercoria* (previously known as *P. copri*), a short-chain fatty acid (acetic acid) producer *Eubacterium siraeum*, probiotic potential candidate *Blautia* and others (Supplemental Table 4) [39–43]. This indicated gut community perturbation in B_12_ deficient condition, where a higher abundance of at least a couple of pathogens and a lower abundance of a few beneficial bacteria (Figure 3D) was observed. Further, the known B_12_ producers among the differentially abundant species were examined: the B_12_ deficient group had about ten known B_12_ producers, while only six were observed in the non-deficient group (Figure 3D).

### Cooperativity-based B_12_ biosynthesis pathway abundance showed no difference in the B_12_ deficient group

Anticipating the incompleteness of the literature information on B_12_ producers, the pathway abundance was further inferred directly from the metagenomic data, and its distribution was compared between the two groups. The pathway abundance in each sample was estimated assuming a cooperative interaction among community members for its biosynthesis. The abundance of all five B_12_ biosynthesis pathways/subpathways (Supplemental Table 5) was found to be statistically similar (*p > 0.05*), indicating no difference between the groups for the given assumption (Figure 4A-C; Supplemental Figure 3A and B).

**Figure 4.**
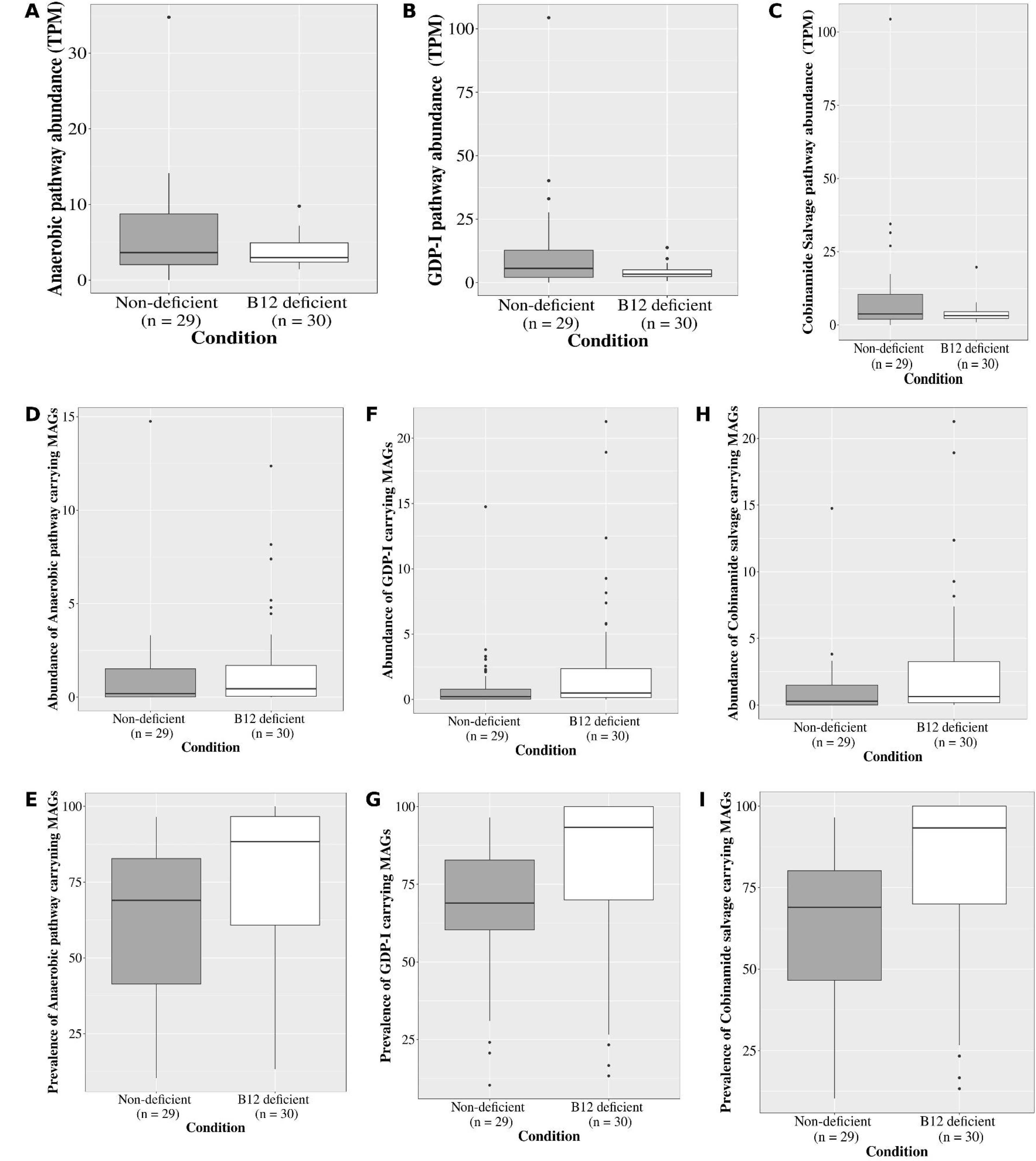
B_12_ Biosynthesis pathway abundance in two groups. **A** anaerobic (*p-value = 0.48*), **B** GDP-I (*p-value = 0.09*), **C** Cobinamide salvage (*p-value = 0.32*). Abundance (median contig coverage) and Prevalence (%) of potential B_12_ producers in two groups. **D** anaerobic pathway abundance (*p-value = 0.303*), **E** prevalence (*p-value = 0.007*), **F** GDP-I pathway abundance (*p-value = 0.01*), **G** prevalence (*p-value = 1.246e-06*), **H** Cobinamide salvage abundance (*p-value = 0.03*), **I** prevalence (*p-value = 2.817e-06*).

### Potential B_12_ producers were generally more prevalent and abundant in the B_12_ deficient group

An alternative to the cooperativity-based B_12_ biosynthesis assumption would be considering the contribution from the MAGs with a complete B_12_ biosynthesis pathway (i.e., the potential B_12_ producers) for abundance estimation. Out of the total of 324 MAGs in the non-deficient group and 367 in the B_12_ deficient group, 135 and 161 were complete, respectively. While the number of MAGs having a particular B_12_ biosynthesis pathway varied between both groups, their proportion was statistically similar (Table 1). Among five pathways, anaerobic, GDP-I, and Cobinamide salvage were present in more than 5 MAGs, so only these were considered for further analysis. The prevalence of the potential B_12_ producers was significantly higher in the B_12_ deficient group for all three pathways; however, their abundance was significantly higher for two of the pathways, with the anaerobic pathway an exception (*p=0.3)* (Figure 4C–I).

### The cumulative abundance of potential B_12_ producers was higher in the B_12_ deficient group

Instead of considering the abundance of potential B_12_ producers for individual pathways, their cumulative abundance in each sample was calculated, and a group-wise comparison was made. The B_12_ deficient group had a much higher cumulative abundance (*p = 7.955e-09*) (Figure 5A). To further establish the association between serum B_12_ level and the cumulative abundance of B_12_ producers, a multiple linear regression analysis was carried out. Results showed a negative association, where with a decline in serum B_12_ levels, the abundance of B_12_ producers significantly increased (*p = <0.001, β_0_ = 6.42, β_1_ = -0.02, Std. Error = 0.006*) (Figure 5B).

**Figure 5.**
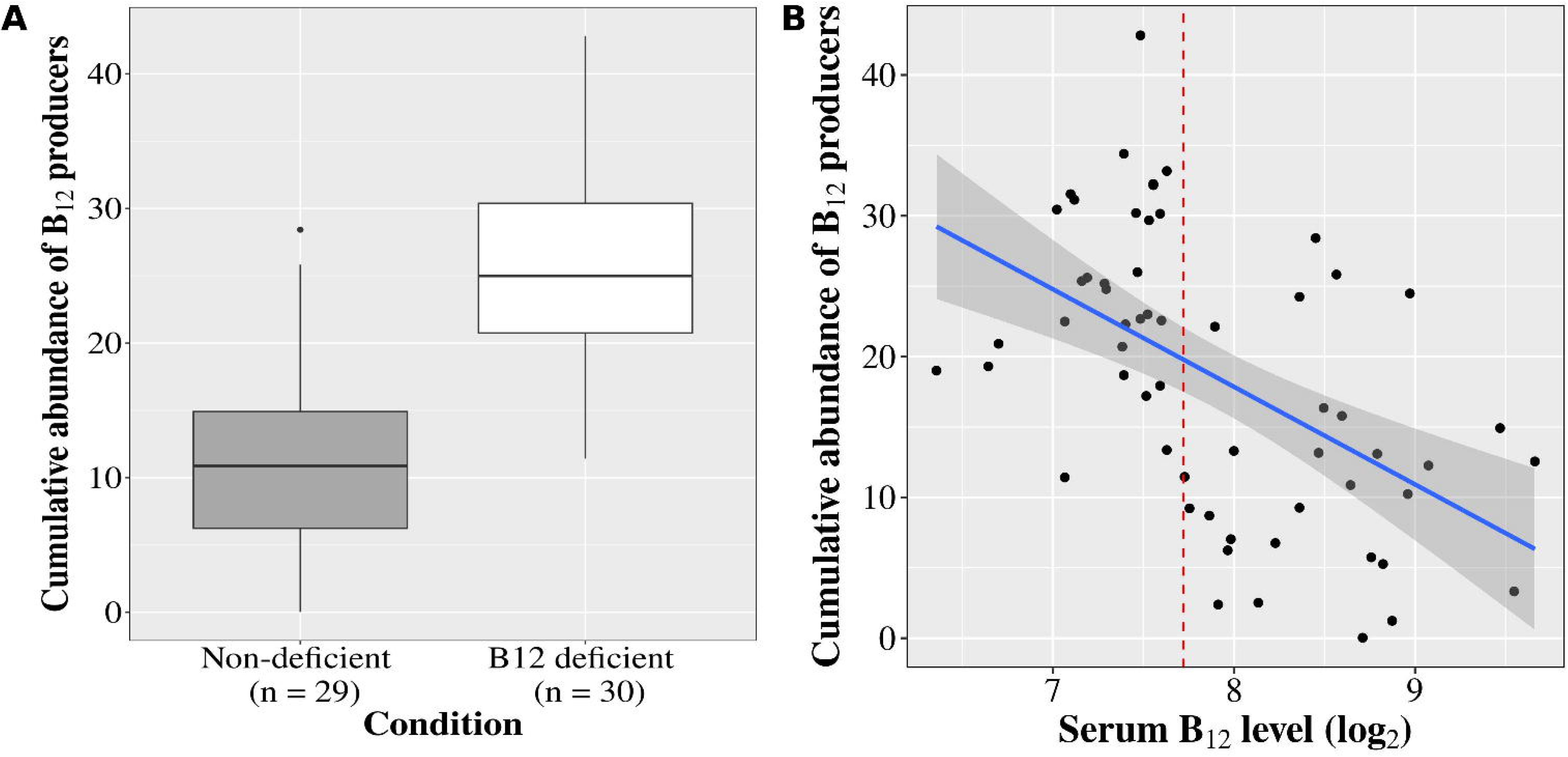
**A** Cumulative abundance of potential B_12_ producers between two groups, **B** Relationship between serum B_12_ level and cumulative abundance of B_12_ producers. The y-axis is the median contig coverage.

Moreover, applying local regression (locally weighted scatterplot smoothing, LOWESS) revealed a refined trend, suggesting that the cumulative abundance of potential B_12_ producers increased non-linearly as the serum B_12_ levels approached the deficient range (Supplemental Figure 4). The majority of the B_12_ producers belonged to the phylum Bacillota, followed by Bacteroidota. 18.4% and 21.21% of them remained unclassified in B_12_ deficient and non-deficient groups, respectively (Supplemental Table 6, Supplemental Figure 5A and B).

## Discussion

To the best of our knowledge, the current study is the first one to directly elucidate the association between serum B_12_ level and gut microbiome at an improved species-level resolution and provide a better understanding than the existing reports. There have been multiple studies in humans and mice to examine the association0 [44–52], but all of them except one were done using 16S rRNA amplicon data and had a resolution at the genus level. The exception was a study on Saudi Arabian females using shotgun metagenome sequencing data; however, their aim was slightly different as they examined the association between Vit. B_12_ status and gut microbiota in the context of obesity [51].

In our study, we observed no significant differences in alpha diversity between the two groups, while a lower beta diversity was noted in the Vit. B_12_ deficient condition. These findings were partially consistent with previous reports. However, the literature evidence on the association between Vit. B_12_ status and gut microbial diversity remains limited. A systematic review on Vit. B_12_ and the gut microbiome summarized that both alpha and beta diversity increased with an increase in Vit. B_12_ levels, as reported in *in-vivo* studies; however, findings from mouse models and observational human studies were heterogeneous [53]. Two of the mouse studies did not show any difference in alpha diversity [46,54], while the third reported lower alpha diversity, but statistical significance was not reported [55]. Similarly, two human studies did not show any difference [44,52], while one reported an increase in alpha diversity with B_12_ intake [45]. Regarding beta diversity, three studies did not show any difference [46,55,56], while one study reported different diversity at the genus level in mice [54]. In humans, two studies showed no difference [44,52], and one showed different beta diversity [45]. Therefore, while majority of the studies have reported similar findings as observed in the current study, others have observed differing outcomes, highlighting variability in the existing literature, possibly due to the difference in sequencing data, sample sizes, experimental variables.

The different microbiome structure was also supported by the differential abundance of a few species in either group (Supplemental Table 4). Along with the enrichment of pathogenic species, the deficient group had a higher abundance of a corrinoid transporter-carrying *Bacteroides thetaiotaomicron,* and several other potential B_12_ producers. A lower relative abundance of the genus *Bacteroides* was reported to be associated with high B_12_ intake in humans [45, 47], and with B_12_ supplementation in mice and in-vitro models [53]. However, this trend was conflicted by another study, which reported no such change in the abundance [44]. *B. thetaiotaomicron*, a mucin generalist, has a crucial role in health and disease pathogenesis [57], and is also known to have multiple copies of corrinoid transporter (BtuFBCD) in its genome, which gives an advantage to this bacterium in distinguishing B_12_ analogs produced by several other members [12, 14, 58]. The higher relative abundance of *B. thetaiotaomicron* in B_12_ deficient groups suggests a scarcity of the B_12_ in the gut. Overall, a higher abundance of genus *Bacteroides* is a characteristic feature of a Vit. B_12_-depleted microenvironment.

Besides, the taxonomic classification of B_12_ producers and their comparison with the available knowledge of producers in the current study also revealed several new species that have biosynthetic potential. For instance, MAGs belonging to species *A. muciniphila, Agathobacter faecis, Agathobacter rectalis, Alistipes senegalensis, Anaerobutyricum hallii, Veillonella atypica etc.,* had B_12_ biosynthesis potential (Supplemental Table 6).

Enrichment of known B_12_ producers and higher prevalence and abundance of potential B_12_ producers in the deficient group, as well as their negative association with serum B_12_ level, indicated that host B_12_ level is linked with the gut microbiome. We hypothesize that this association may reflect a selection pressure on the microbial community to favor those that could grow by synthesizing their own B_12_ over those solely depending on the cobalamin analogs in the lumen. An alternate hypothesis, which may however require more supporting evidence, can be a feedback response possibly originating from the host to the gut microbial community, in a way that enhances microbial B_12_ biosynthesis, so that it could source a fraction of the microbiota-derived B_12_ to meet its DRI and thereby partially prevent acute deficiency. If that being the case, then the gut microbiome would possess buffering ability for homeostasis of some of the essential micronutrients, wherein certain species respond to the micronutrient deficiency in a threshold-dependent manner (at subnormal levels) and undergo microbiome remodeling with an increase in abundance of relevant species for its replenishment. A sustained microbial abundance, along with Vit. B_12_ deficiency of subnormal levels (∼82 pg/ml) indicates a possibility of a threshold-dependent switch-like response (Supplemental Figure 4) that takes care of severe deficiency; however, it is insensitive to achieve clinically sufficient Vit. B_12_ levels. Such a sub-optimal feedback response may be evolutionarily efficient in sparing raw material for *de-novo* Vit. B_12_ biosynthesis under severe micronutrient deficiency in the long term (Supplemental Figure 6).

In our past work, we highlighted the higher abundance of B-vitamin producers and biosynthesis pathways in tribal cohorts as a desirable feature and discussed possible links to different diets, limited access to antibiotics, etc. [10]. Similarly, Das et al. (2019) reported the Chinese population to have higher B-vitamin biosynthesis pathways compared to the other three nationalities [59]. Having observed higher B_12_ producers and biosynthesis pathways in the serum B_12_ deficient group, the previous findings should be re-interpreted based on the hypothesized feedback model. In both previous reports, the B-vitamin status of subjects was unknown, and the groups with higher abundance suggest that the higher abundance could be due to dietary consumption of just adequate or sub-optimal levels of vitamins that are in a range where the switch of the feedback mechanism is active. This indicates that the vitamin levels are partly derived from dietary sources and partly from the microbiome in the healthy host.

### Limitations and future directions

This study was based on case-control design, and hence cause-effect relationship can’t be established. The results were based on a small sample size involving only children. The profiling of other biomarkers of Vit. B_12_ status (holotranscobalamin, methylmalonic acid, and homocysteine) and profiling of biochemical markers to determine the comprehensive health and nutritional status of an individual was not done, which restricted us from taking into account other confounding effects in the analysis. To ascertain the proposed hypothesis and causality of the observed differences, longitudinal/interventional studies required to be done. Further, the microbial signatures identified in our study could also be experimentally investigated in a longitudinal study on an animal model by administering B_12_ deficient and sufficient diets.

## Conclusion

This is the first study to directly elucidate the association between serum B_12_ level and gut microbiome at the species or strain level. The gut community in the B_12_-deficient group was enriched with mucin-degrading bacteria, a few pathogens, and known B_12_-producers. With a higher occurrence of potential B_12_-producers in the deficient group and their negative association with serum B_12_ level, we hypothesize that the gut microbiome responds to the deficiency by selectively promoting the growth of producers, which meets the community’s demand for Vit. B_12_.

## Acknowledgements

We would like to acknowledge the University of Hyderabad’s Institution of Eminence (IoE) for funding our research and the Centre for Modelling, Simulations, and Design (CMSD) for allowing the use of their High-performance Computer facility. Dr Suyash Shrivastava, and Ms Prathysha, Scientists/Project staff from ICMR-National Institute of Research in Tribal Health, Jabalpur, Madhya Pradesh, for preparing the questionnaire; Dr. Devaraj Parasannanavar, Scientist D, ICMR-National Institute of Health, Hyderabad, for his valuable suggestions for grouping samples based on clinical data.

## Authors’ contributions

Designed research: VT, AKV, NC; prepared questionnaire: AKV; collected samples and data: PP, AKV; performed data analysis: NC, VT; performed statistical analysis: NC, PR; Interpreted findings: NC, VT, PR; wrote manuscript: NC, PP, VT, PR; All authors have read and approved the final manuscript.

## Data Availability

The raw data is publicly available at NCBI Sequencing Read Archive under Bioproject ID PRJNA1114651.

## Funding

The project and manpower involved in this research were financially supported by funding from the University of Hyderabad-Institution of Eminence (grant number: UoH-IoE-RC2-21-023) from 30 Aug 2021 to 30th Aug 2024.

### List of abbreviations

p-a dj: Adjusted p-value
CNNS: Comprehensive National Nutrition Survey
IF: Intrinsic factor
MAGs: Metagenome-Assembled Genomes
MCM: Mitochondrial L-Methyl-malonyl-CoA mutase
MetH: Methionine synthase
NRC: Nutrition Rehabilitation Centre
RDI: Recommended dietary intake
Vit. B_12_: Vitamin B_12_

---

**Supplemental Figure 1.** Summary of sampled B_12_ level and fecal microbiome sequencing data. **A** Distribution of serum B_12_ levels in Non-deficient and B_12_ deficient groups. **B** Distribution of average reads per sample before and after filtering.

**Supplemental Figure 2.** Diversity trends. The Box plots represent diversity between Non-deficient and B_12_ deficient groups: **A** Simpson diversity (*p-value = 0.10*), and **B** Robust aitchison (*p-value = 0.07*).

**Supplemental Figure 3. B_12_ Biosynthesis pathway abundance in two groups. A** GDP-II (*p-value = 0.47*), **B** Cobalamin salvage *(p-value = 0.84)*

**Supplemental Figure 4.** Locally weighted scatterplot smoothing (LOWESS) of data in Figure 5B showed that the cumulative abundance of potential B_12_ producers (median contig coverage) increased non-linearly when the serum B_12_ level lies in the deficient range.

**Supplemental Figure 5. Taxonomic classification of potential B_12_ producers at the phylum level in**: A Non-deficient group, **B** B_12_ deficient group.

**Supplemental Figure 6.** The putative Host-microbiome feedback regulation loop **A.** In case of sufficient Vit. B_12_ bioavailability, the serum Vit. B_12_ is maintained above the deficiency threshold, wherein the microbiome shares the Vit. B_12_ from the host with a relatively lesser contribution towards *de-novo* Vit. B_12_ synthesis due to putative inhibitory effect of serum Vit. B_12_ levels on microbiome B_12_ synthesis. **B.** In the case of insufficient Vit. B_12_ uptake or transport (due to lack of intrinsic factor), serum Vit. B_12_ levels fall below a threshold that leads to microbiome remodeling to enhance the abundance of Vit. B_12_ producers and the inhibitory effect on Vit. B_12_ synthesis is relieved so that it is sourced from the microbiome origin to stall the severe Vit. B_12_ deficiency.

**Supplemental** Table 1. Metadata of samples selected for metagenomic analysis.

**Supplemental** Table 2. Age-wise prevalence of B_12_ deficiency in tribal children from Balaghat.

**Supplemental** Table 3. Taxonomic profile of all samples.

**Supplemental** Table 4. Differentially abundant species in B_12_ deficient group with respect to the non-deficient group.

**Supplemental** Table 5. Genes involved in various pathways and subpathways of cobalamin biosynthesis.

**Supplemental** Table 6. Taxonomic classification of vitamin B_12_ pathway carrying MAGs identified in both the groups.

## Reference

1. Roth, JR., Lawrence, JG., Rubenfield, M., Kieffer-Higgins, S., Church, GM. Characterization of the cobalamin (vitamin B_12_) biosynthetic genes of Salmonella typhimurium. Journal of bacteriology. 1993 Jun;175(11):3303–16.

2. Roth, JR., Lawrence, JG., Bobik, TA., Cobalamin (coenzyme B_12_): synthesis and biological significance. Annual review of microbiology. 1996 Oct;50(1):137–81.

3. Allen, RH., Stabler, SP., Savagem DG., Lindenbaum, J., Metabolic abnormalities in cobalamin (vitamin B_12_) and folate deficiency. The FASEB journal. 1993 Nov;7(14):1344–53.

4. Koury, MJ., Ponka, P., New insights into erythropoiesis: the roles of folate, vitamin B _12_, and iron. Annu. Rev. Nutr.. 2004 Jul 14;24(1):105–31.

5. Allen, LH. Causes of vitamin B_12_ and folate deficiency. Food Nutr Bull. 2008 Jun;29(2 Suppl):S20-34; discussion S35-7. doi: 10.1177/15648265080292S105. PMID: 18709879.

6. Green, R., Miller, JW., Vitamin B_12_ deficiency. In Vitamins and hormones 2022 Jan 1 (Vol. 119, pp. 405–439). Academic Press.

7. Stabler, SP., Allen, RH., Vitamin B_12_ deficiency as a worldwide problem. Annu. Rev. Nutr.. 2004 Jul 14;24(1):299–326.

8. National Institute of Health; Office of dietary supplements, https://ods.od.nih.gov/factsheets/VitaminB12-HealthProfessional/ Accessed 30 Oct 2024

9. Rowley, CA., Kendall, MM., To B_12_ or not to B_12_: Five questions on the role of cobalamin in host-microbial interactions. PLoS pathogens. 2019 Jan 3;15(1):e1007479.

10. Chandel, N., Somvanshi, PR., Thakur, V., Characterisation of Indian gut microbiome for B-vitamin production and its comparison with Chinese cohort. British Journal of Nutrition. 2024 Feb;131(4):686–97.

11. Magnúsdóttir, S., Ravcheev, D., de Crécy-Lagard, V., Thiele, I., Systematic genome assessment of B-vitamin biosynthesis suggests co-operation among gut microbes. Frontiers in genetics. 2015 Apr 20;6:148.

12. Degnan, PH., Barry, NA., Mok, KC., Taga, ME., Goodman, AL., Human gut microbes use multiple transporters to distinguish vitamin B_12_ analogs and compete in the gut. Cell Host & Microbe. 2014 Jan 15;15(1):47–57.

13. Degnan, PH., Taga, ME., Goodman, AL., Vitamin B_12_ as a modulator of gut microbial ecology. Cell metabolism. 2014 Nov 4;20(5):769–78.

14. Allen, RH., Stabler, SP., Identification and quantitation of cobalamin and cobalamin analogues in human feces. The American journal of clinical nutrition. 2008 May 1;87(5):1324–35.

15. Comprehensive National Nutrition Survey (2016-2018) https://nhm.gov.in/WriteReadData/l892s/1405796031571201348.pdf Accessed 20 Dec 2024

16. Andrews, S., Babraham Bioinformatics-FastQC a Quality Control Tool for High Throughput Sequence Data. 2010 https://www.bioinformatics.babraham.ac.uk/projects/fastqc (accessed August 2024)

17. Ewels, P., Magnusson, M., Lundin, S., Käller, M., MultiQC: summarize analysis results for multiple tools and samples in a single report. Bioinformatics. 2016 Oct 1;32(19):3047–8.

18. Chen, S., Zhou, Y., Chen, Y., Gu, J., fastp: an ultra-fast all-in-one FASTQ preprocessor. Bioinformatics. 2018 Sep 1;34(17):i884–90.

19. Langmead, B., Salzberg, SL., Fast gapped-read alignment with Bowtie 2. Nature methods. 2012 Apr;9(4):357–9.

20. Li, H., Handsaker, B., Wysoker, A., Fennell, T., Ruan, J., Homer, N., et al. The sequence alignment/map format and SAMtools. bioinformatics. 2009 Aug 15;25(16):2078–9.

21. Blanco-Míguez, A., Beghini, F., Cumbo, F., McIver, LJ., Thompson, KN., Zolfo, M., et al. Extending and improving metagenomic taxonomic profiling with uncharacterized species using MetaPhlAn 4. Nature Biotechnology. 2023 Nov;41(11):1633–44.

22. Dixon, P. VEGAN, a package of R functions for community ecology. Journal of vegetation science. 2003 Dec;14(6):927–30.

23. Li, D., Luo, R., Liu, CM., Leung, CM., Ting, HF., Sadakane, K., et al. MEGAHIT v1. 0: a fast and scalable metagenome assembler driven by advanced methodologies and community practices. Methods. 2016 Jun 1;102:3–11.

24. Seemann, T. Prokka: rapid prokaryotic genome annotation. Bioinformatics. 2014 Jul 15;30(14):2068–9.

25. Li, B., Dewey, CN., RSEM: accurate transcript quantification from RNA-Seq data with or without a reference genome. BMC bioinformatics. 2011 Dec;12:1–6.

26. Caspi, R. Billington, R. Ferrer, L. Foerster, H. Fulcher, CA. Keseler, IM. et al. The MetaCyc database of metabolic pathways and enzymes and the BioCyc collection of pathway/genome databases. Nucleic acids research. 2016 Jan 4;44(D1):D471–80.

27. The National Center for Biotechnology Information. https://www.ncbi.nlm.nih.gov/ Accessed 01 Aug 2024

28. Bairoch, A., Apweiler, R., Wu, CH., Barker, WC., Boeckmann, B., Ferro, S., et al. The universal protein resource (UniProt). Nucleic acids research. 2005 Jan 1;33(suppl_1):D154-9.

29. Emms, DM., Kelly, S., OrthoFinder: phylogenetic orthology inference for comparative genomics. Genome biology. 2019 Dec;20:1–4.

30. Alneberg, J., Bjarnason, BS., de Bruijn, I., Schirmer, M., Quick, J., Ijaz, UZ., et al. CONCOCT: clustering contigs on coverage and composition. arXiv preprint arXiv:1312.4038. 2013 Dec 14.

31. Chklovski, A., Parks, DH., Woodcroft, BJ., Tyson, GW., CheckM2: a rapid, scalable and accurate tool for assessing microbial genome quality using machine learning. Nature Methods. 2023 Aug;20(8):1203–12.

32. Chaumeil, PA., Mussig, AJ., Hugenholtz, P., Parks, DH., GTDB-Tk: a toolkit to classify genomes with the Genome Taxonomy Database.

33. Love, MI., Huber, W., Anders, S., Moderated estimation of fold change and dispersion for RNA-seq data with DESeq2. Genome biology. 2014 Dec;15:1–21.

34. Wexler, AG., Schofield, WB., Degnan, PH., Folta-Stogniew, E., Barry, NA., Goodman, AL., Human gut Bacteroides capture vitamin B_12_ via cell surface-exposed lipoproteins. Elife. 2018 Sep 18;7:e37138.

35. Hiippala, K., Kainulainen, V., Kalliomäki, M., Arkkila, P., Satokari, R., Mucosal prevalence and interactions with the epithelium indicate commensalism of Sutterella spp. Frontiers in microbiology. 2016 Oct 26;7:1706.

36. Muñiz Pedrogo, DA. Chen, J. Hillmann, B. Jeraldo, P. Al-Ghalith, G. Taneja, V., et al. An increased abundance of Clostridiaceae characterizes arthritis in inflammatory bowel disease and rheumatoid arthritis: a cross-sectional study. Inflammatory bowel diseases. 2019 Apr 11;25(5):902–13.

37. Derrien, M., Collado, MC., Ben-Amor, K., Salminen, S., de Vos, WM., The Mucin degrader Akkermansia muciniphila is an abundant resident of the human intestinal tract. Applied and environmental microbiology. 2008 Mar 1;74(5):1646–8.

38. Scheiman, J., Luber, JM., Chavkin, TA., MacDonald, T., Tung, A., Pham, LD., et al. Meta-omics analysis of elite athletes identifies a performance-enhancing microbe that functions via lactate metabolism. Nature medicine. 2019 Jul;25(7):1104–9.

39. Huang, YY., Lu, YH., Liu, XT., Wu, WT., Li, WQ., Lai, SQ., et al. Metabolic properties, functional characteristics, and practical application of Streptococcus thermophilus. Food Reviews International. 2024 Feb 17;40(2):792–813.

40. Precup, G., Vodnar, DC., Gut Prevotella as a possible biomarker of diet and its eubiotic versus dysbiotic roles: a comprehensive literature review. British Journal of Nutrition. 2019 Jul;122(2):131–40.

41. Moore, WC., Johnson, JL., Holdeman, LV., Emendation of Bacteroidaceae and Butyrivibrio and descriptions of Desulfomonas gen. nov. and ten new species in the genera Desulfomonas, Butyrivibrio, Eubacterium, Clostridium, and Ruminococcus. International journal of systematic and evolutionary microbiology. 1976 Apr;26(2):238–52.

42. Liu, X., Mao, B., Gu, J., Wu, J., Cui, S., Wang, G., et al. Blautia—a new functional genus with potential probiotic properties?. Gut microbes. 2021 Jan 1;13(1):1875796.

43. Valentini, L., Pinto, A., Bourdel-Marchasson, I., Ostan, R., Brigidi, P., Turroni, S., et al. Impact of personalized diet and probiotic supplementation on inflammation, nutritional parameters and intestinal microbiota–The “RISTOMED project”: Randomized controlled trial in healthy older people. Clinical nutrition. 2015 Aug 1;34(4):593–602.

44. Boran, P., Baris, HE., Kepenekli, E., Erzik, C., Soysal, A., Dinh, DM., The impact of vitamin B_12_ deficiency on infant gut microbiota. European journal of pediatrics. 2020 Mar;179:385–93.

45. Gurwara, S., Ajami, NJ., Jang, A., Hessel, FC., Chen, L., Plew, S., et al. Dietary nutrients involved in one-carbon metabolism and colonic mucosa-associated gut microbiome in individuals with an endoscopically normal colon. Nutrients. 2019 Mar 13;11(3):613.

46. Lurz, E., Horne, RG., Määttänen, P., Wu, RY., Botts, SR., Li, B., et al. Vitamin B_12_ deficiency alters the gut microbiota in a murine model of colitis. Frontiers in nutrition. 2020 Jun 5;7:83.

47. Carrothers, JM., York, MA., Brooker, SL., Lackey, KA., Williams, JE., Shafii, B., et al. Fecal microbial community structure is stable over time and related to variation in macronutrient and micronutrient intakes in lactating women. The Journal of nutrition. 2015 Oct 1;145(10):2379–88.

48. Forgie, AJ., Pepin, DM., Ju, T., Tollenaar, S., Sergi, CM., Gruenheid, S., Willing, BP., Over supplementation with vitamin B_12_ alters microbe-host interactions in the gut leading to accelerated Citrobacter rodentium colonization and pathogenesis in mice. Microbiome. 2023 Feb 3;11(1):21.

49. Babakobi, MD., Reshef, L., Gihaz, S., Belgorodsky, B., Fishman, A., Bujanover, Y., Effect of maternal diet and milk lipid composition on the infant gut and maternal milk microbiomes. Nutrients 2020; 12: 2539 [Internet].

50. Selma-Royo, M., García-Mantrana, I., Calatayud, M., Parra-Llorca, A., Martínez-Costa, C., Collado, MC., Maternal diet during pregnancy and intestinal markers are associated with early gut microbiota. European journal of nutrition. 2021 Apr;60:1429–42.

51. Al-Musharaf, S., Aljuraiban, GS., Al-Ajllan, L., Al-Khaldi, N., Aljazairy, EA., Hussain, SD, et al. Vitamin B_12_ status and gut microbiota among Saudi females with obesity. Foods. 2022 Dec 11;11(24):4007.

52. Herman, DR., Rhoades, N., Mercado, J., Argueta, P., Lopez, U., Flores, GE., Dietary habits of 2-to 9-year-old American children are associated with gut microbiome composition. Journal of the Academy of Nutrition and Dietetics. 2020 Apr 1;120(4):517–34.

53. Guetterman, HM., Huey, SL., Knight, R, Fox, AM., Mehta, S., Finkelstein, JL., Vitamin B-12 and the gastrointestinal microbiome: a systematic review. Advances in Nutrition. 2022 Mar 1;13(2):530–58.

54. Kelly, CJ., Alexeev, EE., Farb, L., Vickery, TW., Zheng, L., Eric, LC., Kitzenberg, DA., Battista, KD., Kominsky, DJ., Robertson, CE., Frank, DN., Oral vitamin B_12_ supplement is delivered to the distal gut, altering the corrinoid profile and selectively depleting Bacteroides in C57BL/6 mice. Gut Microbes. 2019 Nov 2;10(6):654–62.

55. Zhu, X., Xiang, S., Feng, X., Wang, H., Tian, S., Xu, Y., Shi, L., Yang, L., Li, M., Shen, Y., Chen, J., Impact of cyanocobalamin and methylcobalamin on inflammatory bowel disease and the intestinal microbiota composition. Journal of agricultural and food chemistry. 2018 Dec 20;67(3):916–26.

56. Park, S., Kang, S., Kim, DS., Folate and vitamin B-12 deficiencies additively impaire memory function and disturb the gut microbiota in amyloid-β infused rats. International journal for vitamin and nutrition research. 2019 Dec 16.

57. Wang, C., Zhao, J., Zhang, H., Lee, YK., Zhai, Q., Chen, W., Roles of intestinal bacteroides in human health and diseases. Critical reviews in food science and nutrition. 2021 Nov 25;61(21):3518–36.

58. Desai, MS., Seekatz, AM., Koropatkin, NM., Kamada, N., Hickey, CA., Wolter, M., et al. A dietary fiber-deprived gut microbiota degrades the colonic mucus barrier and enhances pathogen susceptibility. Cell. 2016 Nov 17;167(5):1339–53.

59. Das, P., Babaei, P., Nielsen, J., Metagenomic analysis of microbe-mediated vitamin metabolism in the human gut microbiome. BMC genomics. 2019 Dec;20:1–1.

