## Supplementary Figures for "Inverse association between serum vitamin B_12_ level and abundance of potential B_12_-producing gut microbes in Indian children"

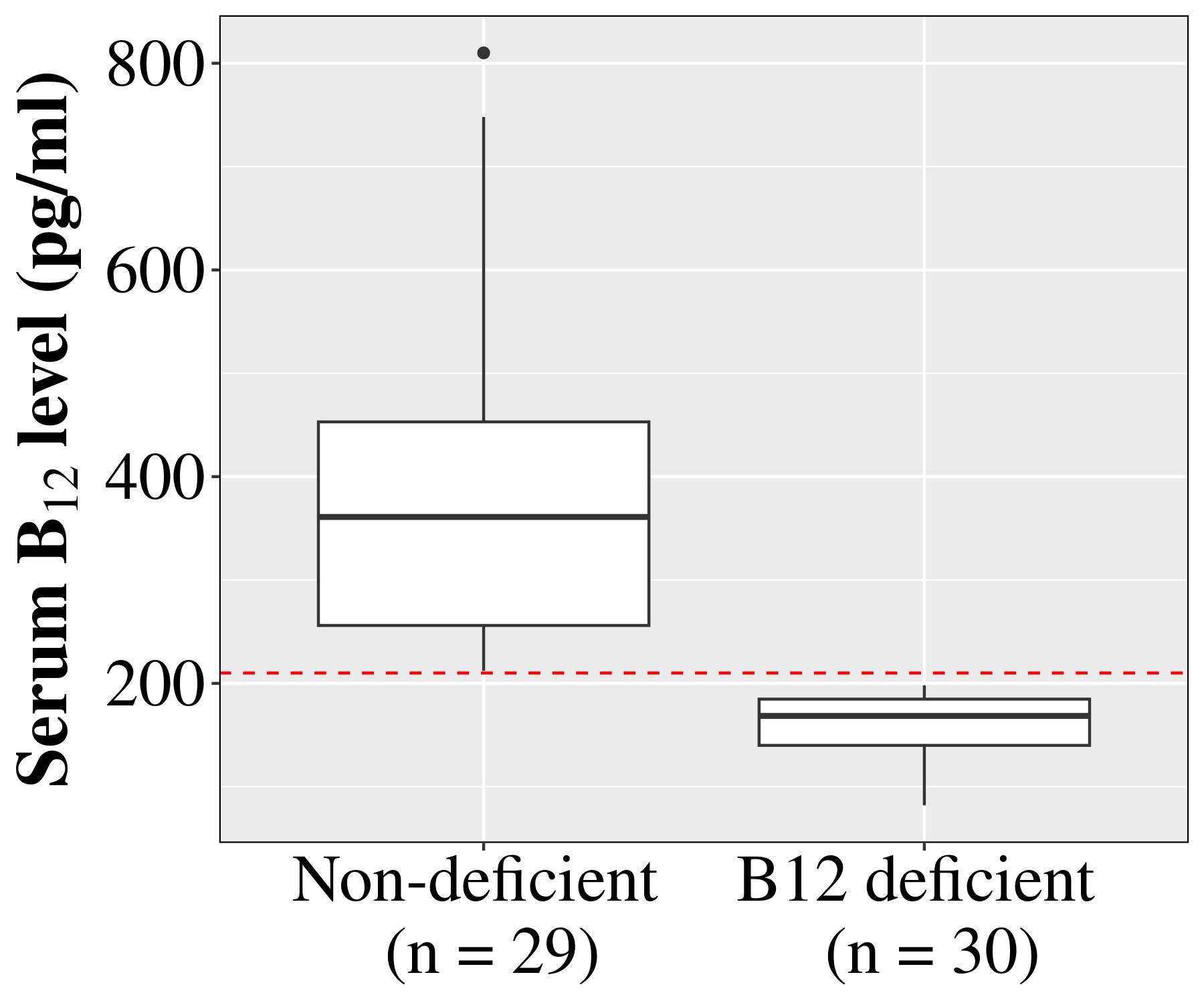

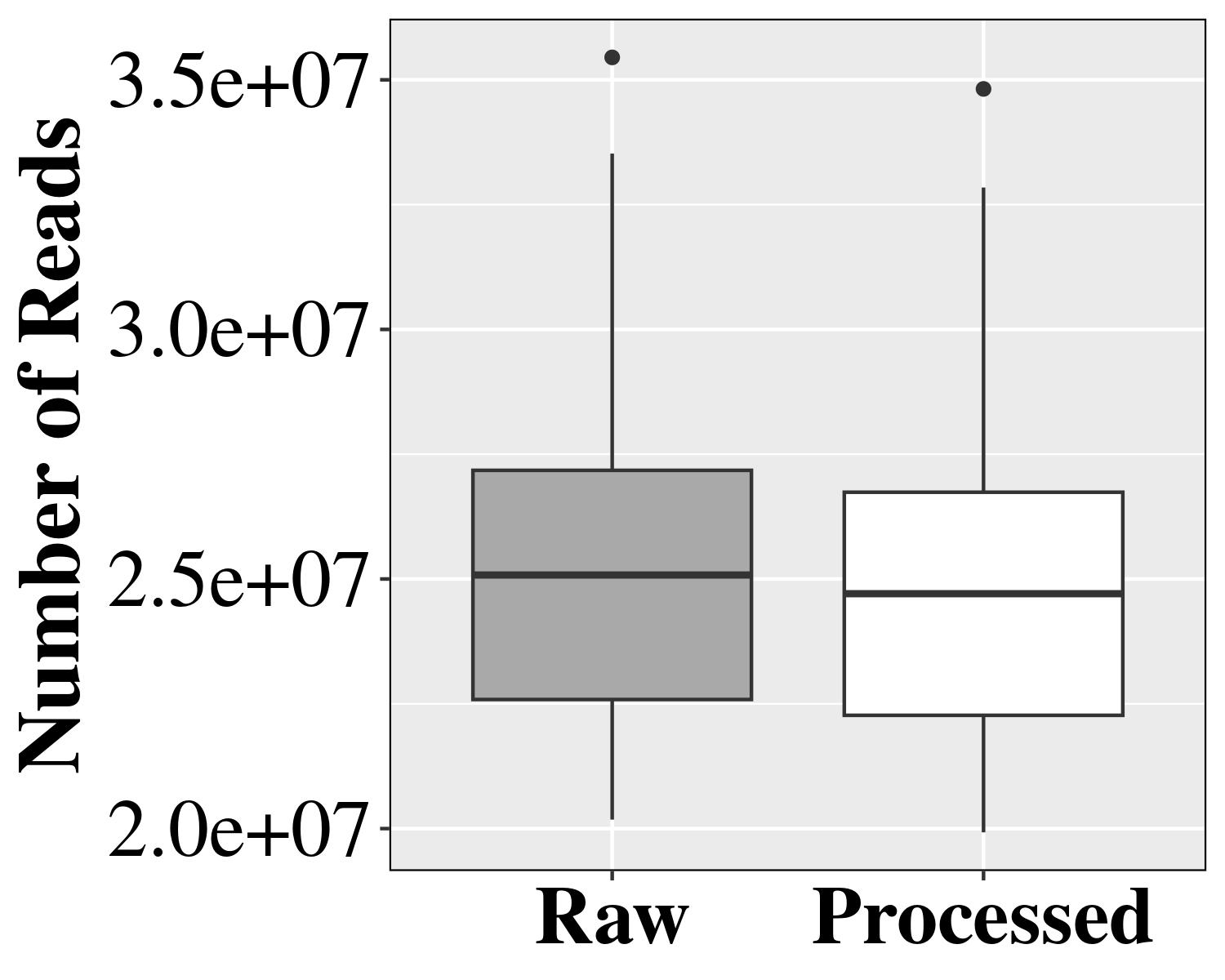

**B**

**A**

**Supplemental Figure 1.** **Summary of sampled B_12_ level and fecal microbiome sequencing data**. **A** Distribution of serum B_12_ levels in Non-deficient and B_12_ deficient groups. **B** Distribution of average reads per sample before and after filtering.

**B**

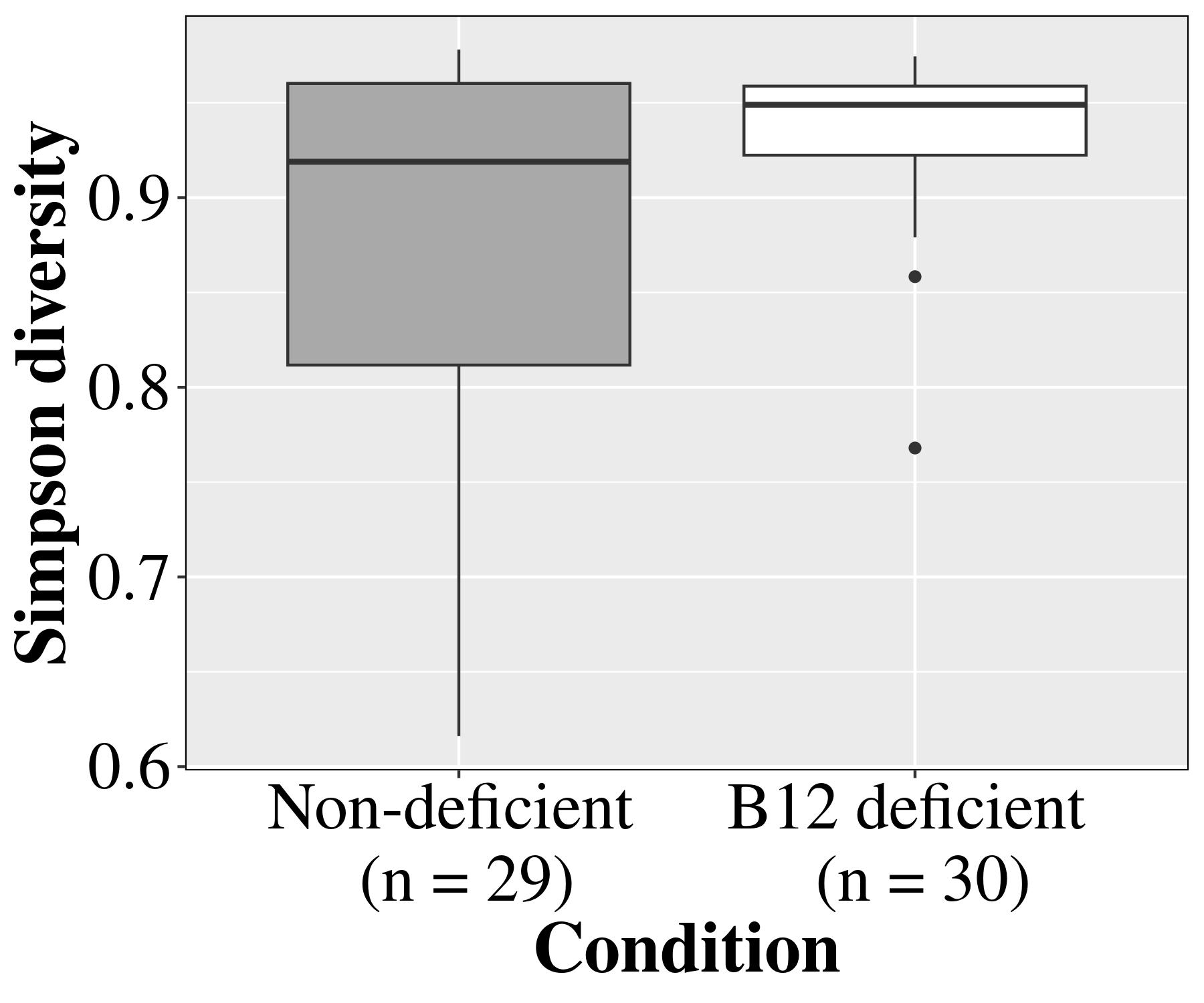

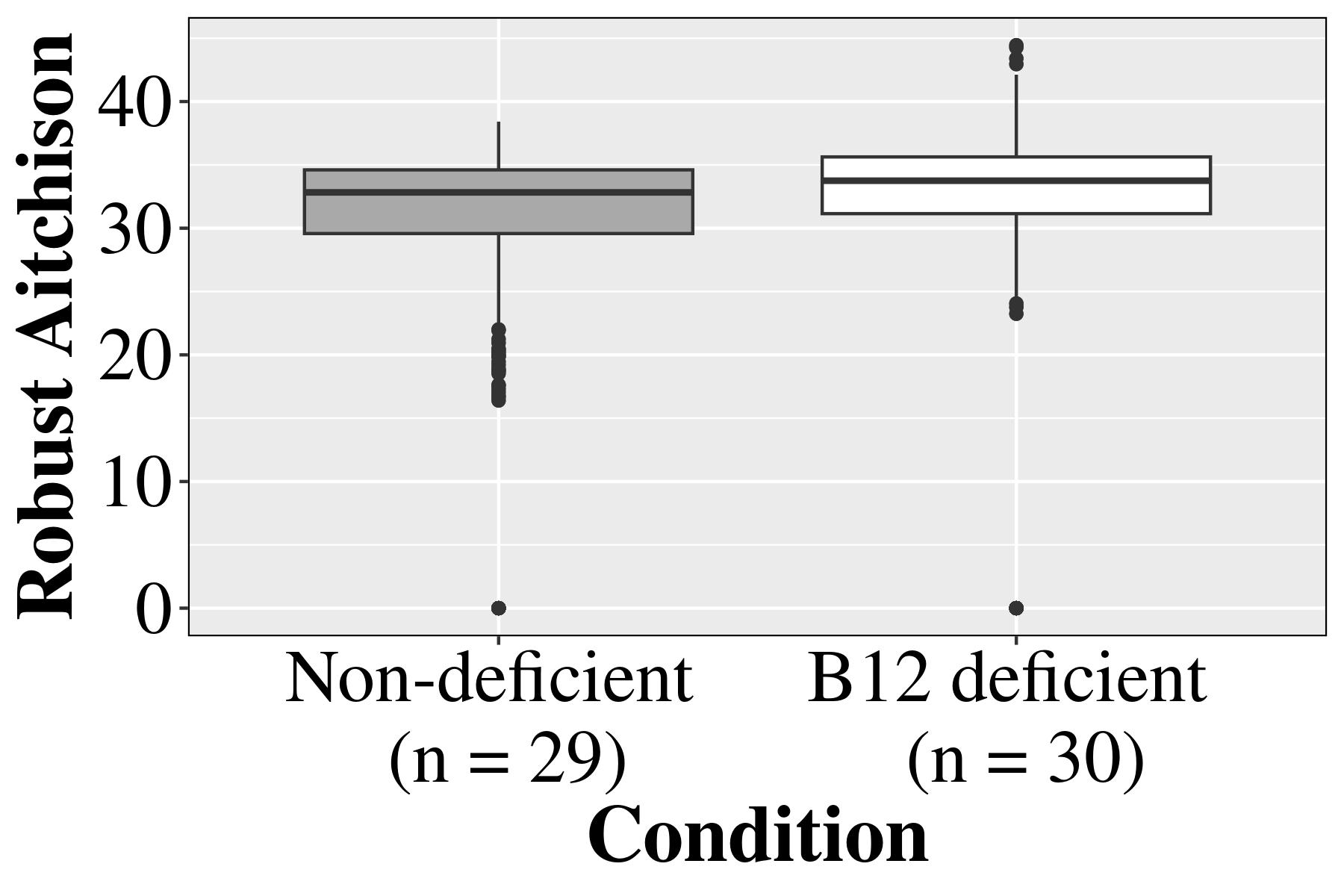

**A**

**Supplemental Figure 2. Diversity trends. The Box plots represent diversity between Non-deficient and B_12_ deficient groups: A** Simpson diversity (*p-value = 0.10*), and **B** Robust aitchison (*p-value = 0.07*).

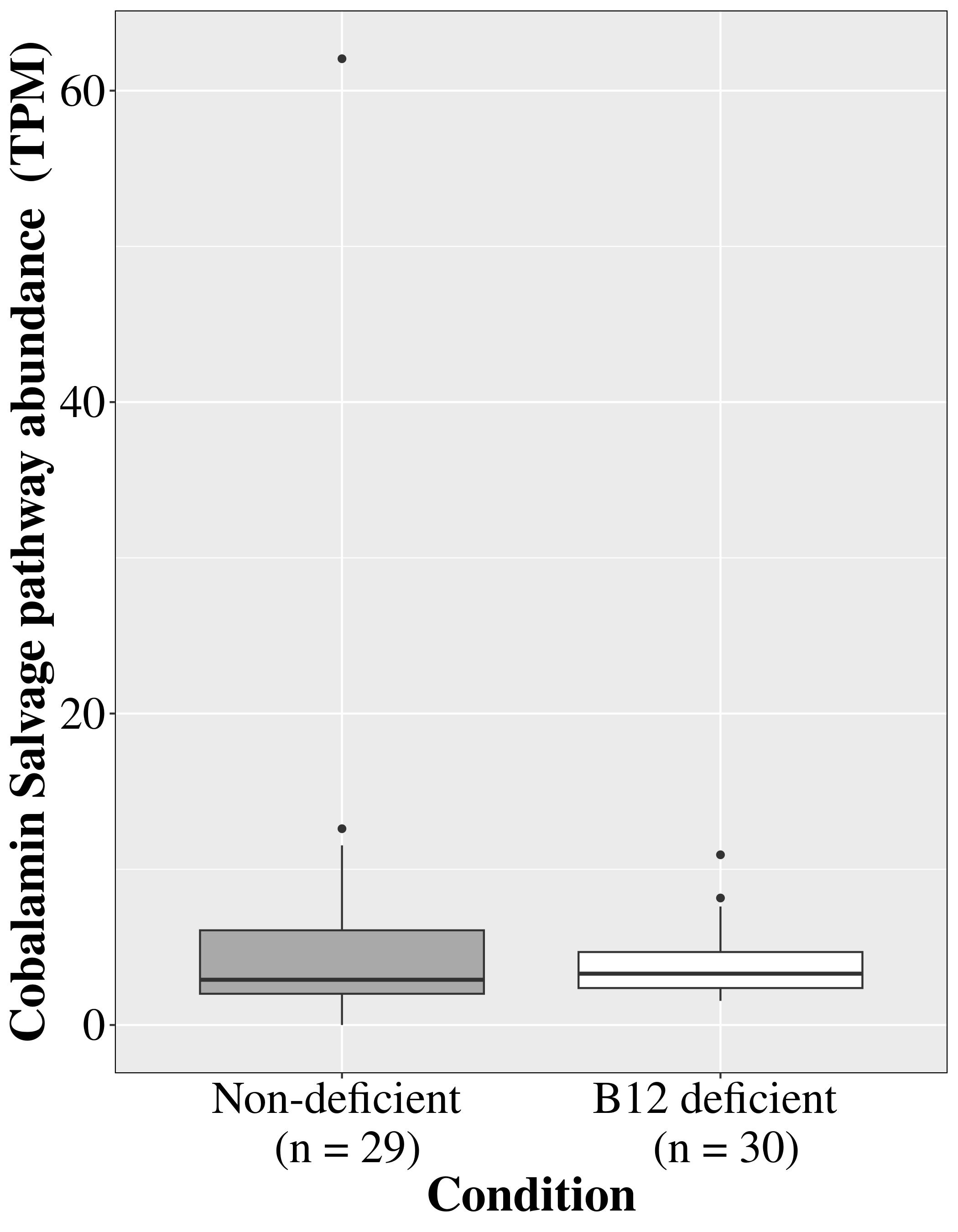

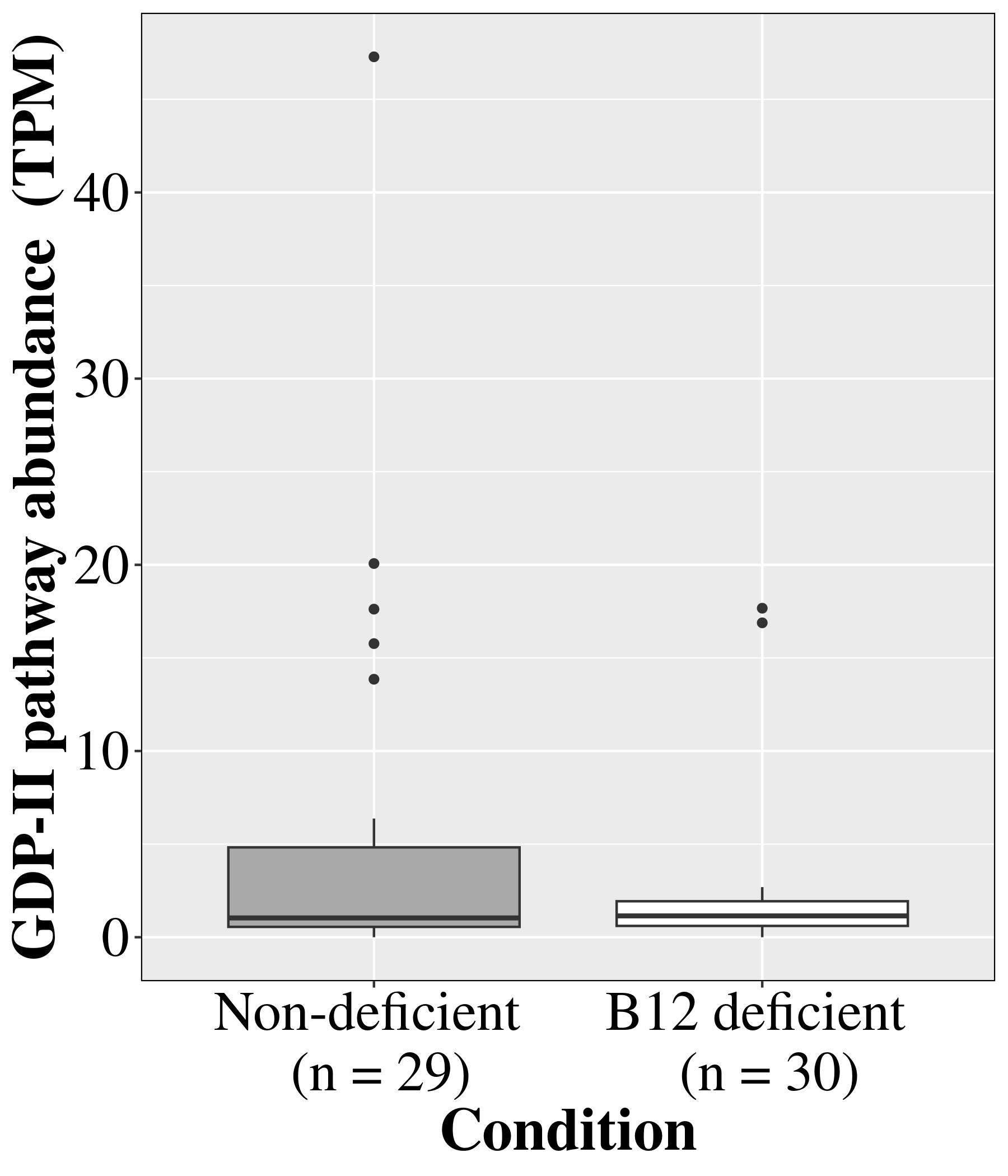

**B**

**A**

**Supplemental Figure 3.** **B_12_ Biosynthesis pathway abundance in two groups.** **A** GDP-II (*p-value = 0.47*), **B** Cobalamin salvage (*p-value = 0.84*)

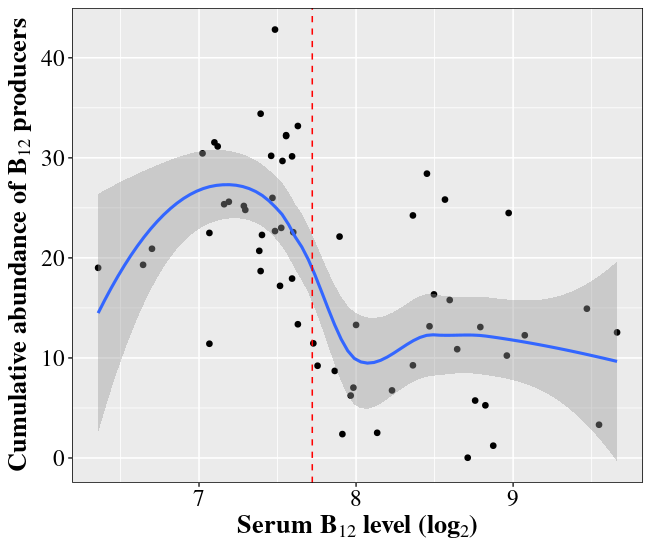

**Supplemental Figure 4.** Locally weighted scatterplot smoothing (LOWESS) of data in Figure 5B showed that cumulative abundance of potential B_12_ producers (median contig coverage) increased non-linearly when the serum B_12_ level lies in the deficient range.

**B**

**A**

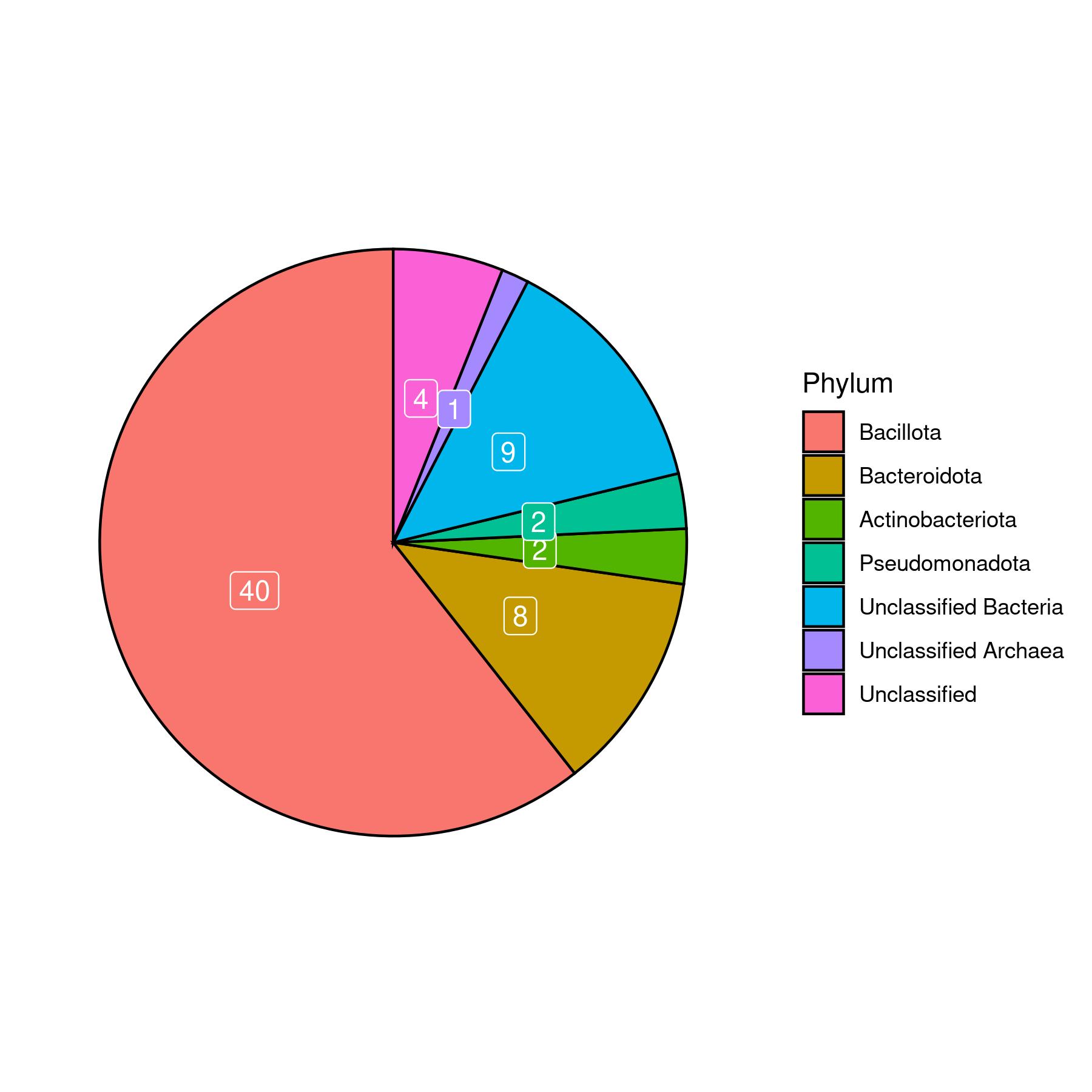

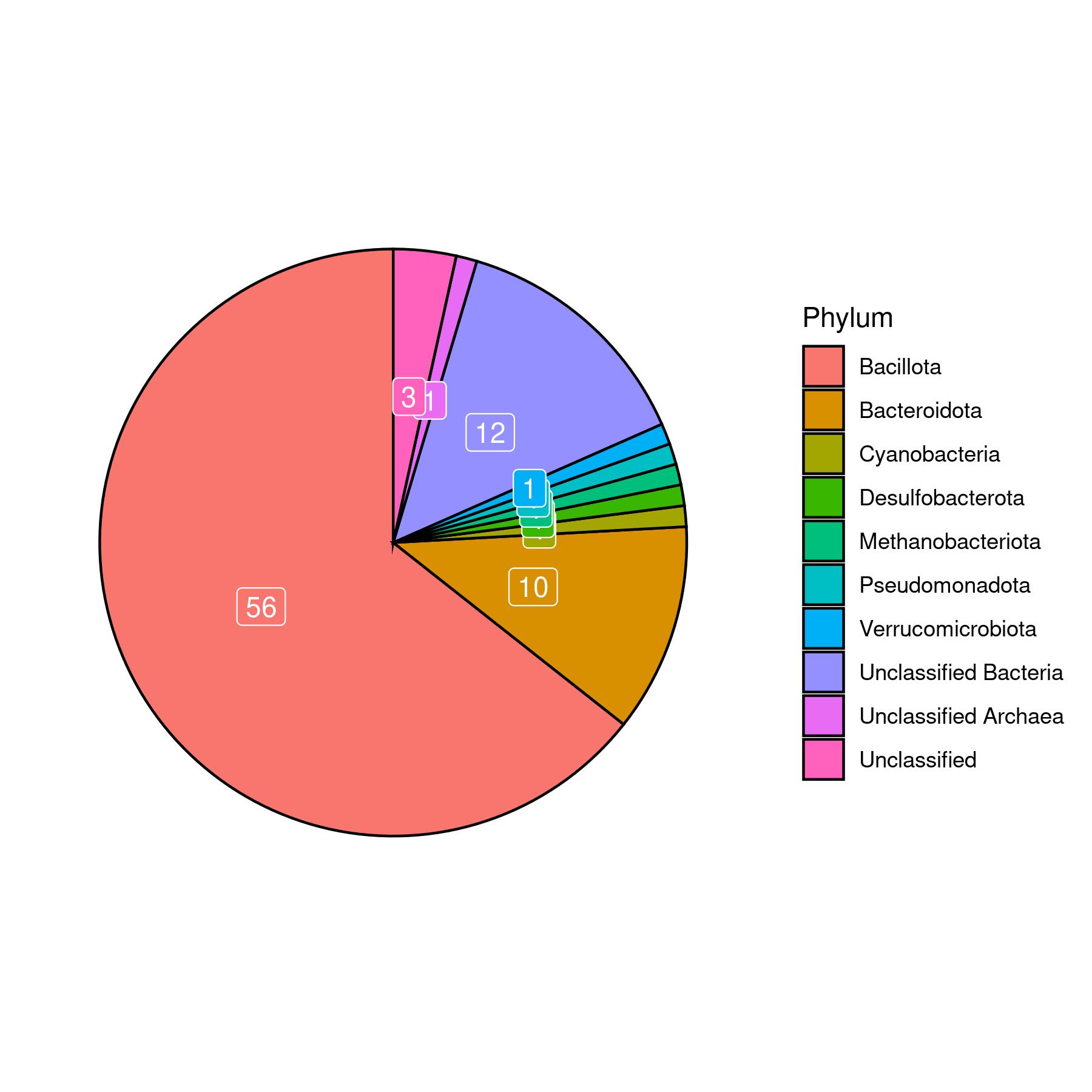

**Supplemental Figure 5.** **Taxonomic classification of potential B_12_ producers at phylum level in**: **A** Non-deficient group, **B** B_12_ deficient group.

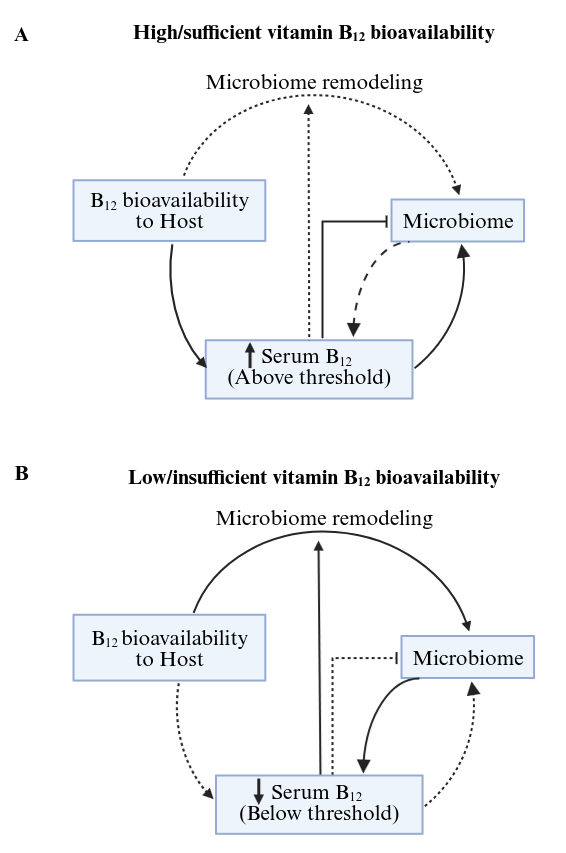

**Supplemental Figure 6.** The putative Host-microbiome feedback regulation loop. **A** In case of sufficient Vit. B_12_ bioavailability, the serum Vit. B_12_ is maintained above the deficiency threshold, wherein the microbiome shares the Vit. B_12_ from the host with a relatively lesser contribution towards *de-novo* Vit. B_12_ synthesis due to putative inhibitory effect of serum Vit. B_12_ levels on microbiome B_12_ synthesis. **B** In the case of insufficient Vit. B_12_ uptake or transport (due to lack of intrinsic factor), serum Vit. B_12_ levels fall below a threshold that leads to microbiome remodeling to enhance the abundance of Vit. B_12_ producers and the inhibitory effect on Vit. B_12_ synthesis is relieved so that it is sourced from the microbiome origin to stall the severe Vit. B_12_ deficiency.
